# Irregular optogenetic stimulation waveforms can induce naturalistic patterns of hippocampal spectral activity

**DOI:** 10.1101/2022.09.21.508935

**Authors:** Eric R. Cole, Thomas E. Eggers, David A. Weiss, Mark J. Connolly, Matthew C. Gombolay, Nealen G. Laxpati, Robert E. Gross

**Affiliations:** Wallace H. Coulter Department of Biomedical Engineering, Georgia Institute of Technology and Emory University, Atlanta, GA, 30332, USA; Department of Neurosurgery, Emory University School of Medicine, Atlanta, GA, 30322, USA; Emory National Primate Research Center, Emory University, Atlanta, GA, 30329, USA; Institute for Robotics and Intelligent Machines, Georgia Institute of Technology, Atlanta, GA, 30332, USA; Department of Neurology, Emory University School of Medicine, Atlanta, GA, 30322, USA

**Author notes:** **Corresponding Author:** Robert E. Gross, MD, PhD.

**Keywords:** medial septum, theta rhythm, brain stimulation, optogenetic, hippocampus

## Abstract

**Introduction:** Brain stimulation is a fundamental and effective therapy for neurological diseases including Parkinson’s disease, essential tremor, and epilepsy. One key challenge in delivering effective brain stimulation is identifying the stimulation parameters, such as the amplitude, frequency, contact configuration, and pulse width, that induce an optimal change in symptoms, behavior, or neural activity. Most clinical and translational studies use constant-frequency pulses of stimulation, but stimulation with irregular pulse patterns or non-pulsatile waveforms might induce unique changes in neural activity that could enable better therapeutic responses. Here, we comprehensively evaluate several optogenetic stimulation waveforms, report their differing effects on hippocampal spectral activity, and compare these induced effects to activity recorded during natural behavior.

**Methods:** Sprague-Dawley rats were prepared for pan-neuronal excitatory optogenetic stimulation of the medial septum (hSyn-ChR2) and 16-channel microelectrode recording in CA1 and CA3 layers of the hippocampus. We performed grid and random sampling of the parameters comprising several stimulation waveforms, including standard pulse, nested pulse, sinusoid, double sinusoid, and Poisson pulse waveforms.

**Results:** We comprehensively report the effects of changing stimulation parameters in these parameter spaces on two key biomarkers of hippocampal function, theta (4-10 Hz) and gamma (32-50 Hz) power. Similarly, robust excitation of hippocampal gamma power was observed across all waveforms, whereas no set of stimulation parameters was sufficient to consistently increase power in the theta band beyond baseline levels of activity (despite the prominent role of the medial septum in pacing hippocampal theta oscillations). Using a manifold learning algorithm to compare high-dimensional neural activity, we show that irregular stimulation patterns produce differing effects with respect to multi-band patterns of activity and can induce activity patterns that more closely resemble activity recorded during natural behavior than conventional parameters.

**Conclusion:** Our counter-intuitive findings – that stimulation of the medial septum ubiquitously does not increase hippocampal theta power, and that different waveforms have similar effects on single power bands – contradict recent trends in brain stimulation research, necessitating greater caution and fewer mechanistic assumptions as to how a given stimulation target or waveform will modulate a neurophysiological biomarker of disease. We also reveal that irregular stimulation patterns can have biomimetic utility, promoting their exploration in medical applications where inducing a particular activity pattern can have therapeutic benefit. Last, we demonstrate a scalable data-driven analysis strategy that can make the discovery of such physiologically informed temporal stimulation patterns more empirically tractable in translational settings.

## 1. Introduction

Brain stimulation is a fundamental tool for understanding the mechanisms of neural circuits and treating neurologic diseases such as Parkinson’s disease, epilepsy, and more.^1-3^ In most of these applications, brain stimulation is conventionally delivered as a train of electrical pulses at a fixed frequency. However, irregular temporal waveforms of stimulation have proven capable of driving more varied patterns of neural activity. This property could lead to greater therapeutic benefit in multiple neuropsychiatric applications^4^ – for example, achieving longer-lasting tremor reduction with reduced energy cost for treating Parkinson’s disease.^5, 6^ However, it is unknown how complex waveforms can modulate neural activity or which patterns to test due to fundamental difficulties in brain stimulation: it is challenging to predict how the brain will respond to different stimulation parameters before directly testing them. Furthermore, it is intractable to naively try every possible stimulation configuration as there are millions of settings available with modern clinical devices and infinitely many in the space of all possible temporal waveforms.^7, 8^ To effectively use irregular temporal stimulation patterns for precision control of neural circuits and symptoms, we need scalable. data-driven strategies to study the effects of massive stimulation parameter spaces on neural activity and identify novel temporal patterns that may be useful for any given application.

In this study, we use optogenetic stimulation of the medial septum and its effect on hippocampal local field potential activity as a model system to study the effects of irregular brain stimulation waveforms and parameters. Optogenetics, the use of light-activated ion channels to excite or inhibit specific populations of neurons, holds several advantages over traditional electrical stimulation for this study: its effects feature high signal-to-noise ratio properties, are replicable within and across subjects, are robust to changes over time, and allow for complete visibility of electrically recorded neural activity during stimulation (as compared to electrical stimulation, which induces artifacts that obscure measured neural activity)^9^. These features make optogenetic stimulation an attractive model to prototype advanced algorithmic strategies to improve and study the effects of brain stimulation, enabling their translation to the improvement of electrical stimulation in more challenging therapeutic domains. The medial septum is a well-characterized pacemaker region that robustly controls oscillations throughout the hippocampal formation, providing an effective model to investigate how variability in stimulation parameters affects control of neural activity.^10^ In addition, a better understanding of how various parameters of medial septum stimulation can control hippocampal oscillations could improve its therapeutic potential as a treatment for epilepsy.^11^

In this study, we test five different classes of optogenetic stimulation waveforms in the medial septum, comprehensively sweep their parameters, and measure their effects on hippocampal activity. First, we characterize the effects of varying the stimulation waveform on single biomarkers with well-characterized roles in hippocampal function – theta power and low gamma power – and find significant but modest differences between the effects of different temporal waveforms on individual frequency bands. We then demonstrate a strategy using dimensionality reduction to show that meaningful differences between stimulation waveforms arise when studying their effects across a broader frequency spectrum of hippocampal local field potential activity. We then compare the induced activity patterns to “natural” neural activity recorded during several behavior paradigms. We find that the activity patterns induced by standard constant-frequency pulsatile stimulation are predominantly artificial (i.e., mostly different from endogenously recorded neural activity), whereas the range of activity patterns induced by all other evaluated parameter spaces aligned more closely with neural activity observed during behavior. These results reveal the benefit of exploring novel temporal stimulation patterns for precise neural control and demonstrate an effective data-driven strategy to discover novel biomimetic patterns of stimulation, which can be broadly used to improve other therapeutic and scientific uses of neuromodulation.

## 2. Methods

### 2.1 Surgical protocol

A total of four adult male Sprague-Dawley rats (200-250 g) from Charles River Laboratories (Wilmington, MA, USA) were used for this study. Each stage of analysis uses a within-subject control over an average of 5500 stimulation trials performed per subject to derive the results. All animals were maintained within a 12-hour light/dark cycle vivarium with *ad libitum* access to food and water. All procedures were conducted in accordance with Emory University’s Institute for Animal Care and Use Committee.

For each subject, two surgical procedures under anesthesia (1.5%–4% inhaled isoflurane) were conducted as previously described.^12^ First, the viral vector AAV5-hSynapsin-Channelrhodopsin2-eYFP was injected into the medial septum at a 20 degree angle to the dorsal-ventral axis (0.40 mm anterior, 2.12 mm lateral at the 20◦ angle from bregma, 5.80 mm ventral to pia along the rotated axis). A volume of 1.8 μl containing 1012 particles/mL was injected with a rate of 0.35 μl/min using a pulled-glass pipette attached to a stereotactically mounted injector (Nanoject, Drummond Scientific Co., Broomall, PA, USA). Once the pipette was withdrawn, the scalp was stapled closed, and Meloxicam was administered as an analgesic (3–5 mg/kg).

A second survival surgery was conducted after two weeks, allowing time for recovery and optogenetic channel expression. A 16-channel multielectrode array (MEA; Tucker Davis Technologies (TDT), Alachua, FL., USA) with a 1-mm depth offset between rows (designed to target hippocampal CA3 and CA1 pyramidal cell layers) was implanted in the right dorsal hippocampus (centered at 3.50 mm posterior and 2.80 mm lateral to bregma). The MEA was lowered ventrally into the brain until pyramidal single unit activity was observed from both the CA1 and CA3 regions. The ferrule was then driven into the reopened original injection craniectomy at a 20-degree angle to the dorsal-ventral axis to approximately 5.1 mm from the pia along the rotated axis. Correct ferrule depth was determined by applying a 17 Hz, 10 ms, 50 mW/mm^2^ stimulation for 10 seconds to verify a change in hippocampal activity. Five 2-mm stainless steel screws were mounted on the skull for electrode’s ground and reference wires as well as for the structural support. Finally, the craniotomy was sealed with dental acrylic to secure the electrode and the ferrule in place. Figure 1 shows a schematic of fiber and multielectrode array placement.

**Figure 1:**
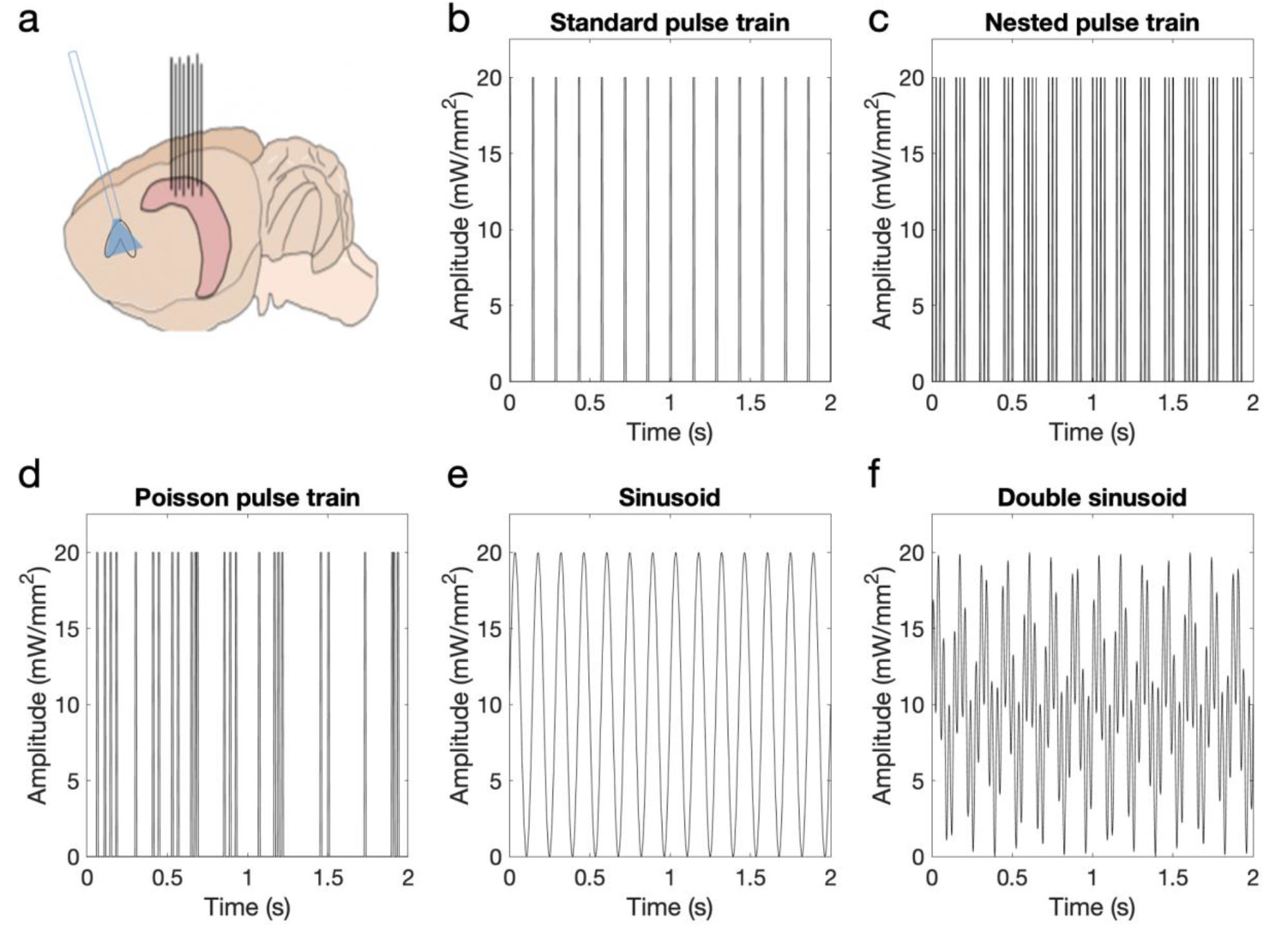
Methods for optogenetic stimulation of the medial septum. a) Schematic of an implanted optical fiber in the medial septum (blue) and 16-channel microelectrode recording from the hippocampus (pink) in the rat brain. b-e) Depiction of optogenetic stimulation waveforms and parameters. b) Standard pulse train waveform with parameters: amplitude = 20 mW/mm^2^, frequency = 7 Hz, pulse width = 5 ms. c) Nested pulse train waveform with parameters: amplitude = 20 mW/mm^2^, train frequency = 7 Hz, pulse frequency = 40 Hz, train width = 80 ms, pulse width = 5 ms. d) Poisson pulse train waveform with parameters: amplitude = 20 mW/mm^2^, frequency = 7 Hz, pulse width = 5 ms. e) Sinusoid waveform with parameters: amplitude = 20 mW/mm^2^, frequency = 7 Hz. f) Double sinusoid waveform with parameters: amplitude = 20 mW/mm^2^, ratio = 0.5, first frequency = 7 Hz, second frequency = 30 Hz.

### 2.2 Stimulation Parameter Search Experiments

After recovering from the second survival surgery, subjects underwent experiments to comprehensively evaluate parameters of different optogenetic stimulation waveforms (shown in Fig. 1 b-f). Throughout this text, we will use the term “waveform” to denote the class of waveform being used (i.e., standard, sine, nested pulse, or double sinusoid), whereas the term “parameter space” will refer to the set of all possible parameter combinations for a given waveform class within the boundaries described below.

1. Standard pulse train (Fig. 1b): Pulsatile time series of light defined by an amplitude (10-50 mW/mm^2^), frequency (5-42 Hz), and width (2-10 ms) of rectangular pulses.
2. Sinusoid (Fig. 1c): Sinusoidal time series defined by an amplitude (10-50 mW/mm^2^) and frequency (2-50 Hz).
3. Poisson pulse train (Fig. 1d): Time series of rectangular pulses where the time interval between pulses is randomly drawn from a Poisson distribution. Parameters include amplitude (10-50 mW/mm^2^), center frequency of the Poisson distribution (5-42 Hz), and pulse width (2-10 ms). A refractory period was used to ensure that the time between pulses was at least 1 ms.
4. Nested pulse train (Fig. 1e): A time series of rectangular pulses defined by amplitude (10-50 mW/mm2), pulse frequency (20-100 Hz) and pulse width (2-5 ms), mediated by a slower on/off rectangular “train” cycle defined by a train frequency (2-12 Hz) and train width (20-80 ms). Note that, for a certain subspace of the parameter space, the parameters may produce either a standard pulse waveform (if the train width is low enough to only fit one pulse in its duration for the given pulse frequency) or other irregularities if the train width and frequency are high enough that adjacent trains overlap in time. During an experiment, any parameter set that met these conditions was discarded and new parameters were sampled before we generated and applied the waveform.
5. Double sinusoid (Fig. 1f): A time series of two superimposed sinusoids, defined by the amplitude (10-50 mW/mm^2^) of the overall signal, two sinusoid frequency values (2-50 Hz), and the relative ratio of each sinusoid’s magnitude vs the signal amplitude (0-1). The signal is created by generating and summating two sinusoids with relative amplitudes determined by the proportion parameter, both at 0 phase, then scaling the resultant signal such that the minimum value is 0 and maximum value is the amplitude.

The frequency values were chosen based on the time constant of ChR2, which has activation kinetics that saturate at frequencies greater than 50 Hz^13^. Amplitude values were chosen based on previous findings that amplitudes higher than 50 mW/mm^2^ in the medial septum saturate the hippocampal response and produce no greater change in gamma power.^14, 15^ Combinations of parameters within these spaces were comprehensively applied during experiments using two sampling strategies: grid search, where parameters are selected according to a predetermined grid; and random search, where all parameter values are drawn from a uniform distribution where the boundaries described above provide the minimum and maximum values. Grid sampling allows us to quantify the variability across repeated samples of a single parameter combination throughout the parameter space, whereas random sampling ensures that values in between the arbitrarily spaced grid points are tested at a smaller resolution as well.

Such experiments were performed during two different behavioral conditions: open field exploratory behavior (an enriched environment derived from a spatial object recognition memory task) and quiescent behavior (a small, opaque black enclosure). These two behaviors were chosen to provide two complementary states of hippocampal activity: elevated theta oscillations during spatial navigation, and depressed theta oscillations during quiescence.^16^ The total number of trials performed during each experiment was adjusted based on the dimensionality of the parameter space: 2 parameters = 600, 3 parameters = 750, 4 or 5 parameters = 1750 (values are the total number of trials performed per subject and are presented as totals over several days of experimentation). Individual trials were performed in 10-second blocks, consisting of 5 seconds for pre-stimulation recording/washout and a 5-second duration waveform at the selected stimulation parameters. Throughout the experiment, hippocampal local field potentials were recorded from the hippocampus using an RZ2 BioAmp Processor and a PZ2 pre-amplifier (Tucker Davis Technologies (TDT), Alachua, FL, USA). Signals were recorded at a sampling rate of 24414 Hz, and then down sampled to 2000 Hz before feature extraction and analysis.

### 2.3 Signal Processing and Feature Calculation

For each trial, the power spectral density (PSD) was computed separately for the 5 second pre-stimulation and during-stimulation LFP recordings and averaged across channels. Channels and individual trials that were manually identified to have poor signal quality due to excess movement artifact or faulty or disconnected wiring and recording equipment during each experiment were excluded from the analysis. The PSD was computed using the multi-taper power spectral density method from the Chronux toolbox^17^, using a time-bandwidth product of 3 with 5 tapers, and extracting the power signal between 0-100 Hz. Individual power features corresponding to canonical neural bands were extracted by integrating the signal in theta (4-10 Hz) and low gamma (32-50 Hz) ranges.

### 2.4 Modeling Effects of Parameters on Single-Band Features

#### 2.4.1 Gaussian process modeling

Gaussian process (GP) regression models were constructed using the MATLAB Gaussian Processes for Machine Learning Toolbox v4.2.^18^ Cross validation was performed to choose the optimal kernel configuration for the Gaussian process regression. Two mean functions (i.e., constant, affine) and seven covariance functions (i.e., Matern, periodic, rational quadratic, squared exponential, Gabor, linear and polynomial) were tested. Kernels with the lowest normalized mean squared error across the four animals was chosen for each feature, which was the affine/Matern kernel for gamma and theta power. Statistical outliers which were more than 3 standard deviations from the GP mean prediction were removed and hyperparameters were optimized within the GP framework. These models used the stimulation parameters as inputs and predicted the expected percent increase in power for gamma and theta. Power increase estimates were normalized by the baseline power (averaged across all stimulation-off epochs in a given experiment session), so that the models predicted the percent change in power.

### 2.5 Dimensionality Reduction Analysis

To derive the neural latent space, the power spectral density was computed as described above for all 5-second stimulation trials in the range of 0-50 Hz, then log-scaled. This was to ensure that both low and high-frequency regions of the power spectrum were balanced in magnitude and dimensionality, so that the dimensionality reduction methods used did not overweight or neglect one part of the frequency domain when deriving the neural latent space. The resulting dimensionality of the input space was 164 frequency values. We then applied three different dimensionality reduction methods to reduce the post-processed data to a two-dimensional representation:

#### 2.5.1: Uniform Manifold Approximation and Projection (UMAP)

Uniform Manifold Approximation and Projection (UMAP) is a nonlinear manifold learning technique that falls under the class of k-neighbor based graph learning algorithms.^19^ Such an algorithm proceeds in two phases: first, the data is used to construct a k-neighbor weighted graph, a network representation where a “weight” is used to quantify the similarity between each data point and its *k* nearest neighbors. Second, a lower-dimensional set of points is created that preserves the structure of this graph (e.g., the weights between each data point and its neighbors are equivalent to those in the original k-neighbor weighted graph). For most of our results, we focus on the latent space derivation provided by UMAP as it is particularly well-suited to balancing the influence of both local and global structure in data (as opposed to emphasizing or overfitting to one of these properties).^20^ We provide a shortened description of the UMAP algorithm:

1a) Compute a distance metric for each pair of points in the high-dimensional data set (in our implementation, the Euclidean distance), and find each data point’s respective set of k nearest neighbors that have the smallest pairwise distance.
1b) A fuzzy k-neighbor local metric ball is used to create an individual weighted graph of local similarity for each data point *j* with its k neighbors. First, a ball (assigned a radius of 1) is fitted to each individual data point *j* such that the ball’s radius stretches to the closest of point *j*’s neighbors. For the rest of point *j*’s k neighbors (each of which has a distance greater than the unit ball radius), an exponential distribution is used to quantify the weight of their connection (theoretically, the “probability” that the graph connection between j and k exists) with the centered data point, scaled based on the local density of the data set.
1c) The local similarity graphs for each point *j* are used to construct the global k-neighbor weighted graph for the whole data set, where the weight between two points j and k is averaged based on their two respective local similarity graphs by the formula *w*_*j,k*_ = *w*_*j*→*k*_ + *w*_*k*→*j*_ − (*w*_*j*→*k*_ ∗ *w*_*k*→*j*_).
2a) A force directed graph layout algorithm is used to minimize the distance between a new k-neighbor weighted graph, *H*, made of two-dimensional points (y_1_, y_2_) and the high-dimensional k-neighbor weighted graph. The graph, *H*, is first initialized to an arbitrary layout based on the spectral layout of the high-dimensional graph (or, this could be done randomly or using a different heuristic).
2b) This graph configuration then provides the starting point for a non-convex optimization problem, solved using force directed graph layout, where the objective is to find a new graph, *H*’, that minimizes the edge-wise cross-entropy distance between it and the original graph derived from the high-dimensional data. Force directed graph layout solves this optimization problem by iteratively updating the graph H using attractive and repulsive forces: an attractive force is applied to pull each pair of points closer together, and then a repulsive force is applied to push adjacent points farther apart, at each iteration to update the structure of graph H towards a smaller cross-entropy metric. The magnitude of these forces are slowly decreased throughout the optimization process to assure convergence to a local minimum.

The design of UMAP is strongly grounded in mathematical theory for the purpose of learning a manifold that balances local and global structure in the original data; for a thorough justification as to why this is the case, see the original paper.^19^

#### 2.5.2: t-Distributed Stochastic Neighbor Embedding (t-SNE)

t-distributed stochastic neighbor embedding (t-SNE) is another nonlinear manifold learning method that falls under the class of k-neighbor based graph learning algorithms.^21^ This algorithm works by computing pairwise distances between the high-dimensional data points, modeling the set of distances as a probability distribution, then creating an equivalent set of lower-dimensional points that minimizes the Kullback-Leibler divergence between it and the original probability distribution. In practice, t-SNE is thought to primarily preserve local structure rather than global structure present in the high-dimensional data; i.e., clusters of points very close to each other in the latent space would mostly accurately represent distances between these points in the original data, but distances between point clusters that are far apart in the latent space are less likely to reflect the original distance values.^22^

#### 2.5.3: Principal Component Analysis (PCA)

We also performed principal component analysis (PCA), which provides a linear method to derive the neural latent space based on its covariance structure.^23^ PCA creates orthogonal bases, or linear combinations of input features, which attempt to maximally preserve covariance in the data. This method necessarily captures global structure present in the high-dimensional data, at the cost of discarding nonlinearity and local structure.

### 2.6 Analysis in the Neural Latent Space

Each of the described dimensionality reduction algorithms was used to derive a two-dimensional representation of the high-dimensional data collected during stimulation for each subject. After applying each method, outlier trials were excluded by computing the low-dimensional Euclidean distance between all pairs of points, then discarding trials exceeding a mean of three standard deviations of distance away from all other points in the low-dimensional space. To avoid the confounding scenario where low-amplitude stimulation parameters may appear similarly to behavioral trials by simply producing no or little effect on neural activity, all stimulation trials with low-amplitude parameters (less than 30 mW/mm^2^) were excluded from analysis before deriving the neural latent space.

#### Boundary identification

In the neural latent space, a boundary was identified for each parameter space to delineate the range of neural activity patterns that was produced by the given waveform. First, coordinates in the space were normalized such that the minimum and maximum X and Y coordinates fall within the range [-1,1]. Then, to identify the boundaries associated with a given stimulation parameter space, kernel density estimation was used to fit a 2D nonparametric probability distribution where the probability is proportional to the density of points at a given point in space in each group.^24^ We used a two-dimensional symmetric Gaussian kernel with standard deviation of 0.05. A boundary for each group was identified as the region of space where p(x,y) > 0.01. This boundary identification method was used to identify boundaries for data from each parameter space behavioral data, as well as the total data set collected for a given subject (after outlier exclusion as described above).

#### Quantifying boundary area

For each region of interest, the normalized area of each region of interest was computed by dividing the area of each individual boundary by the area of the total boundary capturing all data points for a given subject (after outlier exclusion). The Sorenson-DICE coefficient was used to quantify the overlap between different regions of interest as shown in the equation below, where |X| and |Y| represent the area in pixels of regions X and Y, respectively.^25^

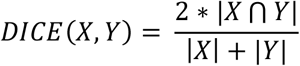

This metric is equal to 1 when both boundaries are exactly overlapping, 0 when they do not overlap, and is low when the regions overlap partially but have different sizes and/or shapes.

## 3. Results

### 3.1 Parameter Effects on Hippocampal Low Gamma Power

Hippocampal LFP recordings were collected from four subjects during optogenetic stimulation of the medial septum delivered at a comprehensive set of stimulation parameters and waveforms. Figure 2a shows the LFP recording (averaged across hippocampal channels and repeated trials) during different stimulation waveforms at an equivalent stimulation frequency of 35 Hz (the double sine parameters were excluded from this stage of analysis due to the lack of a central frequency parameter; trials at a pulse frequency of 35 Hz were used for nested pulse parameters). The sine and standard waveforms demonstrated coherent high-frequency activity that is higher in magnitude than baseline activity and aligned to the onset of stimulation, while the nested pulse and Poisson patterns did not.

**Figure 2:**
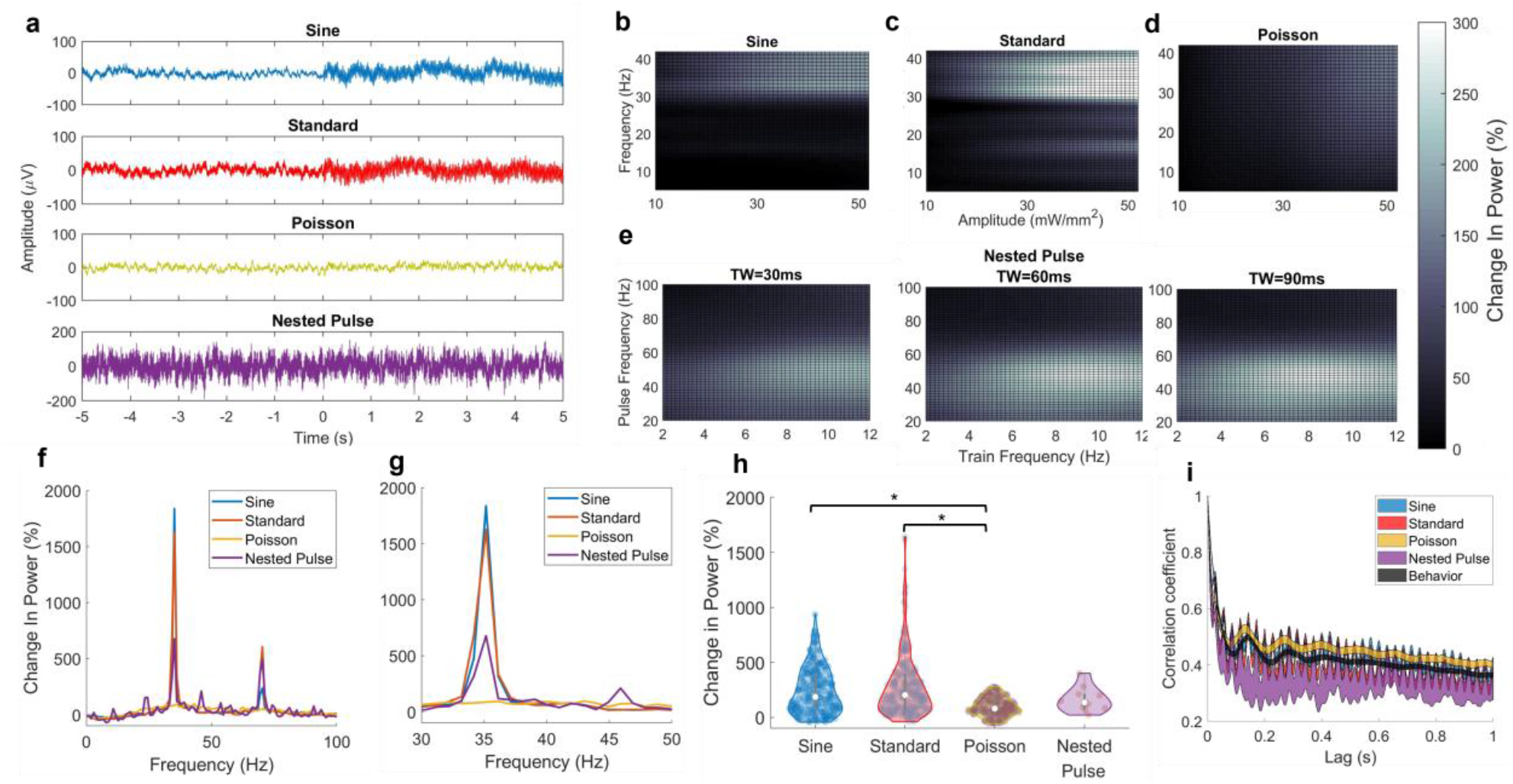
Effect of optogenetic stimulation waveforms on low gamma power. a) Averaged LFP during 5 seconds of 35 Hz stimulation parameters for various waveforms (0 seconds indicates start time of stimulation; -5 through 0 seconds indicates baseline activity). b) Mean surface of Gaussian process regression model predicting the low gamma power percent change vs. baseline as a function of standard pulse parameters. c) Equivalent surface plot for sinusoid parameters. d) Equivalent surface plot for Poisson pulse parameters. e) Equivalent surface plots for nested pulse train parameters, corresponding to three values of the train width: left = 30 ms, middle = 60 ms, and right = 90 ms. f-g) Modulation profile showing the bandpower percent change from baseline throughout the power spectral density. Left: 0-100 Hz; right: 30-50 Hz. h) Change in gamma power relative to baseline for all stimulation waveforms. Sine and standard induced a significantly greater gamma power increase than Poisson, with no other significant differences among waveforms. i) LFP autocorrelation during stimulation for all waveforms at 35 Hz frequency.

For both the sine and standard pulse spaces (Fig. 2b-c), the parameters observed to maximize gamma power corresponded to a maximal optical amplitude (50 I) and a pulse frequency near 40 Hz, whereas lower frequency and amplitude yielded a decreased power change (consistent with our prior observations using this model).^15, 26^ The Poisson pulse train parameters (Fig. 2d) demonstrated similar gamma-maximizing parameters as observed in the standard pulse space, but with a broadening effect of the frequency parameter – a wider range of frequency values was able to produce a lower-amplitude increase in gamma power as compared to the standard pulse space, where a steeper gradient is observed in the regression model surface. In the nested pulse parameter space (Fig. 2e), gamma power remained most sensitive to the optical amplitude and a pulse frequency in the gamma range, with other parameters providing a more modest contribution: a high train frequency was necessary for a maximum response but increasing the train width also increased the gamma response despite lower train frequencies. In the frequency domain, 35 Hz stimulation was accompanied by a peak in 35 Hz power and a harmonic at 70 Hz which were most prominent for standard and sine parameters. Poisson stimulation produced a broadband increase in power centered at 35 Hz, and nested pulse stimulation produced a less sharp 35 Hz peak with several smaller peaks throughout the power spectrum (Fig. 2f-g).

We compared the effects of these stimulation patterns by selecting the subset of trials corresponding to 35 Hz frequency and maximum amplitude (the gamma-maximizing combination for each parameter space): the Poisson pulse parameters yielded a smaller gamma power increase than other parameters, with a statistically significant difference in the Poisson waveform compared to standard and sine stimulation. Distributions were compared with the Mann-Whitney U test with Bonferroni correction for multiple comparisons, using a = 0.05 as the significance level. In an autocorrelation analysis (Fig. 2i), both standard and sine waveforms featured a prominent correlation at 35 Hz which was greatly reduced in the Poisson waveform. The average autocorrelation value across all time lags was higher for Poisson and sine patterns than the other waveforms.

### 3.2 Parameter Effects on Hippocampal Theta Power

Figure 3a shows averaged local field potential recordings for different stimulation waveforms at a 7 Hz stimulation frequency. Consistent with the results for 35-Hz parameters in section 3.1, the standard and sine waveforms evoke a coherent high-magnitude periodic waveform whereas no change was observed during Poisson stimulation. The nested pulse pattern produced a response that is salient but less pronounced than was observed with the other waveforms (trials at a train frequency of 7 Hz were used for nested pulse parameters).

**Figure 3:**
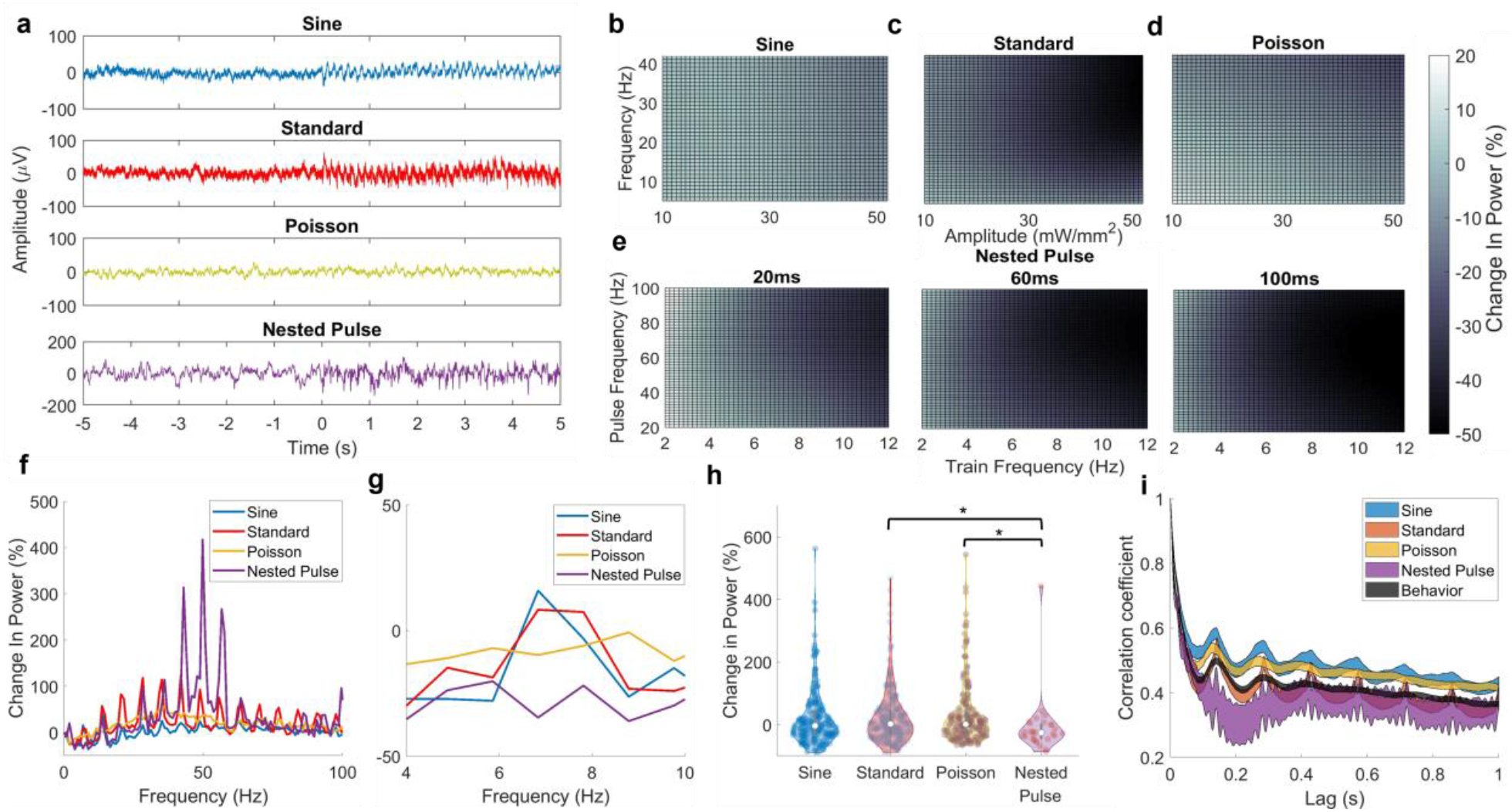
Effect of optogenetic stimulation waveforms on theta power. a) Averaged LFP during 5 seconds of 35 Hz stimulation parameters for various waveforms (0 seconds indicates start time of stimulation; -5 through 0 seconds indicates baseline activity). b) Mean surface of Gaussian process regression model predicting the theta power percent change vs. baseline as a function of standard pulse parameters. c) Equivalent surface plot for sinusoid parameters. d) Equivalent surface plot for Poisson pulse parameters. e) Equivalent surface plots for nested pulse train parameters, corresponding to three values of the train width: left = 30 ms, middle = 60 ms, and right = 90 ms. f-g) Modulation profile showing the bandpower percent change from baseline throughout the power spectral density. Left: 0-100 Hz; right: 4-10 Hz. h) Equivalent frequency and amplitude parameters across waveforms produce no increase in theta power; a significant difference was detected between the nested pulse and standard/Poisson (p<0.05). i) LFP autocorrelation during stimulation for all waveforms at 7 Hz frequency.

Gaussian process regression model predictions are again visualized in Figure 3b-e to demonstrate the mean theta power change vs. stimulation parameters for each space. In general, theta power regression models featured a critically lower sensitivity to the effect of stimulation, with large portions of the parameter space producing an approximate zero-percent change in theta power relative to baseline. The effects of changing stimulation parameters, while apparent, are less pronounced than was observed with gamma power. Sinusoid amplitude and frequency parameters (Fig. 3b) showed negligible sensitivity in the mean response. In the standard pulse space (Fig. 3c), the theta power response was diminished at high amplitudes and frequencies outside of the theta range. Similar to the sine waveform, the theta power response to Poisson pulse (Fig. 3d) parameters was largely insensitive to changes in parameters. In the nested pulse train parameter space (Fig. 3e), a decrease in theta power was observed at higher train frequency values with a limited effect of other parameters. In the frequency domain, harmonic peaks of 7 Hz were observed for multiple waveforms and were weakest for the sine and Poisson waveforms; nested pulse parameters also produced a strong harmonic associated with the pulse frequency (50 Hz). Sine and standard waveforms produced a small increase in power above baseline at exactly 7 Hz but not in the broader range of the theta band (Fig. 3f-g). The effects of these parameter spaces were compared by selecting the subset of trials corresponding to 7Hz and maximum amplitude: no waveform was able to consistently increase theta power beyond its baseline value. The nested pulse waveform was found to produce a significantly greater decrease in theta power compared to standard and Poisson stimulation (p<0.05). All stimulation patterns displayed a similar 7 Hz autocorrelation profile as was observed in the behavioral data, aside from the nested pulse pattern which featured a prominent 50 Hz oscillatory pattern (associated with the pulse frequency) (Fig. 3i). The standard pulse pattern displayed sharp peaks at 7 Hz lag spacings which were not observed in the sine and Poisson waveforms.

Based on the findings that stimulation has a different ability to entrain power in low (theta) and high-frequency (gamma) bands, we examined the effect of all stimulation frequencies on power within the stimulated frequency band (Figure 4). The magnitude of the power entrainment effect monotonically increased with the stimulation frequency. On average, entrainment above baseline power in the stimulated band did not occur below the 10-15 Hz range, and robust power increases were observed above 20 Hz. This trend was consistent across different parameter spaces, but the magnitude of this trend was strongest for the standard pulse parameters and weakest for the Poisson pattern (Fig. 4c). However, the monotonic nature of oscillation entrainment vs. frequency did not substantially deviate for the different stimulation waveforms, aside from the size of the effect. Analysis of the autocorrelational structure in the induced time series showed that the coherence of an oscillation driven at the stimulation frequency did not substantially deviate from the coherence observed at the same frequency during behavior (Fig. 4a-c) – that is, the influence of frequency on the shape and temporal structure of the induced oscillation also did not change based on the waveform.

**Figure 4:**
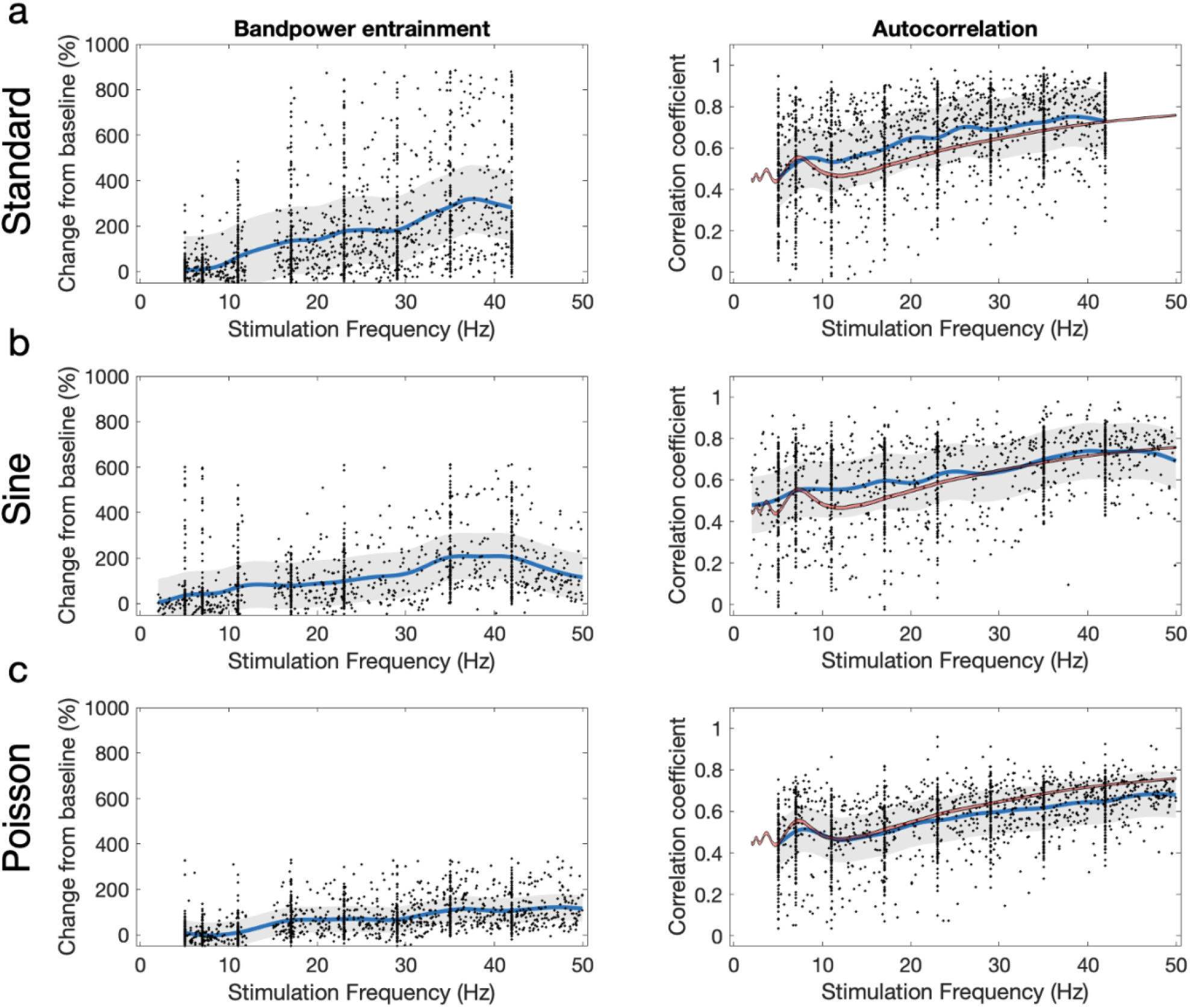
Effectiveness in entraining oscillations vs. stimulation frequency. a-c) Change in power vs. pre-stimulation baseline power (left) and autocorrelation coefficient (right) at the stimulated frequency during stimulation for three parameter spaces (top: standard; middle: sine; bottom: Poisson). (For example, the y-value shown at a stimulation frequency of 35 Hz is the percent change in 35 Hz power, or the autocorrelation coefficient of the time series at a 35-Hz time lag.) Blue lines and gray shading respectively indicate the mean and standard deviation of a Gaussian process regression model fit to the raw data (black points). Red line: average autocorrelation coefficient during sham trials.

### 3.3 Irregular Optogenetic Stimulation Waveforms Can Induce Naturalistic Multi-Band Patterns of Activity

We then investigated the effects of these stimulation parameter spaces by performing dimensionality reduction on the PSD for data during all the stimulation trials in this study, in addition to data recorded during quiescent and exploratory natural behaviors (during which no stimulation was applied). This algorithmic process produces a “neural latent space” representation of the raw PSD data: a two-dimensional set of coordinates where each data point has an equivalent one-to-one match with a data point in the original data set, and the distance relationship between each pair of high-dimensional points is approximately preserved in the two-dimensional representation. Visualization of the neural latent space revealed that individual parameter spaces produce representations that are partially overlapping, but with distinct and separable components (a representative example is shown in Fig. 4a; Fig. S1 shows all representations derived for multiple combinations of subject and reduction algorithm). Notably, the behavioral data organically produced two distinct clusters in the neural latent space, which is consistent with the two behavioral conditions used in the experiment design (visualizable in Fig. 4d and 4e). Figure 4b shows the calculated boundaries for each parameter space (see Methods 2.5.4), and figure 4c shows the normalized area which provides a quantification of the “potential” of each parameter space – the range of different neural activity patterns that could be induced by the given set of stimulation waveforms. All parameter spaces had a greater normalized area than the behavioral data, which had a mean normalized area of 0.28. The standard pulse train space displayed the highest normalized area of 0.70, which was greater than the nested pulse train area and Poisson pulse train area. This would mean that, despite their complexity and higher dimensionality, the size of unconventional parameter spaces (double sine, nested pulse train, and Poisson pulse train) did not equivalently correspond to greater flexibility or a larger range of neural activity patterns that can be induced in the neural latent space.

Similarly, we investigated the extent to which the neural latent space representations produced by these parameter spaces are similar, in comparison to other parameter spaces and behavioral conditions for each animal. Despite occupying a high area, the standard pulse space was found to mainly occupy regions of the neural latent space that are separate from the behavioral region (Fig. 4d), whereas other parameter spaces such as the nested pulse train were found to have greater overlap with the behavioral region (Fig. 4e; Fig. S2 and S3 respectively show the comparison between stimulation and behavior, and between parameter spaces, for all parameter spaces and subjects). A DICE score was used to quantify the overlap between the boundaries corresponding to each parameter space (Fig. 4f). In comparison to activity observed during behavior, the standard pulse train space was found to have the smallest DICE score, whereas all other parameter spaces featured greater overlap with the behavioral data. This result was consistent across different algorithmic methods used to derive the neural latent space (Fig. S4), and therefore was not a byproduct of the specific machine learning strategy used. To avoid the confound that this effect could be produced by low-amplitude stimulation parameters that simply had no effect on neural activity, all stimulation trials with low-amplitude parameters (less than 30 mW/mm^2^) were excluded from analysis before deriving the neural latent space. This concern is also ameliorated by the observation of robust low-gamma power entrainment in each subject (Fig. 2).

**Figure 4:**
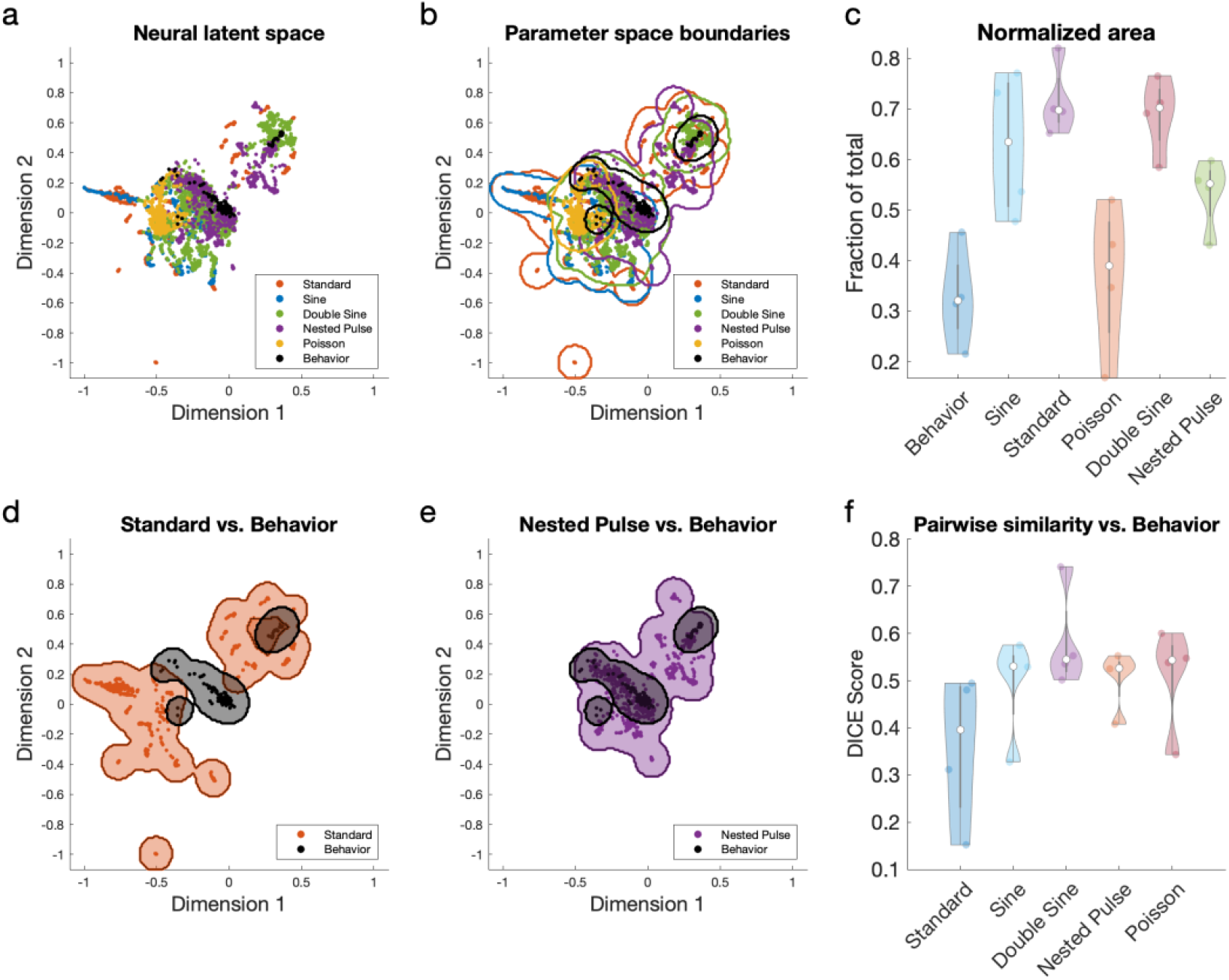
Irregular parameter spaces can induce multi-band activity patterns that more closely resemble behavioral neural activity. a) Representative neural latent space representation via the UMAP algorithm of PSD during all stimulation parameters tested (example shown for one subject that demonstrated the highest-magnitude power response to optogenetic stimulation). b) Representative boundaries computed for individual parameter spaces in the UMAP-derived latent space. c) Normalized area occupied by each parameter space boundary in the latent space (N = 4 subjects). The waveforms are ordered by the dimensionality of their parameter spaces, from smallest (N=2) to greatest (N=5). d) Comparison of boundaries between standard pulse and behavior parameter spaces for the same subject as in a and b. e) Comparison of boundaries between nested pulse and behavior parameter spaces. f) Quantification of the degree to which the boundary of each parameter space overlaps with the boundary of behavior data (N = 4 subjects). Low-amplitude stimulation trials (less than 30 mW/mm^2^) were excluded from this analysis.

Finally, we inspected the neural latent space to find the properties of stimulation parameters and neural activity that underlie these observations of similarity between groups. We manually identified coordinates of interest in the neural latent space – finding the 50 nearest neighbors of the selected point – and then computed the mean PSD and identified the stimulation waveforms and parameters that were associated with the given neural pattern (Fig. 5). We first chose a coordinate inside one cluster of behavioral activity (Fig. 5a), which showed elevated activity within the theta band and the first harmonic of the theta band. The associated stimulation waveforms were predominantly low train frequency nested pulse parameters and double sine parameters containing a mixture of low (theta) and medium-range (beta) frequencies. Another behavioral cluster (Fig. 5b) showed a lower theta peak with no harmonics, which was again accompanied by double sine and nested pulse parameters at a mixture of low (theta-range) and higher (gamma-range) frequencies. An “artificial” pattern that did not overlap with these clusters (Fig. 1c) showed increased broadband activity centered near 30 Hz, which was mainly produced by Poisson parameters. Another “artificial” pattern (Fig. 1d) showed a pattern with multiple sharp harmonics in the PSD, which was produced by 23 Hz sinusoidal parameters, standard parameters at both 23 Hz and 11 Hz frequencies, and one double sine parameter with beta and gamma-range frequencies.

**Figure 5:**
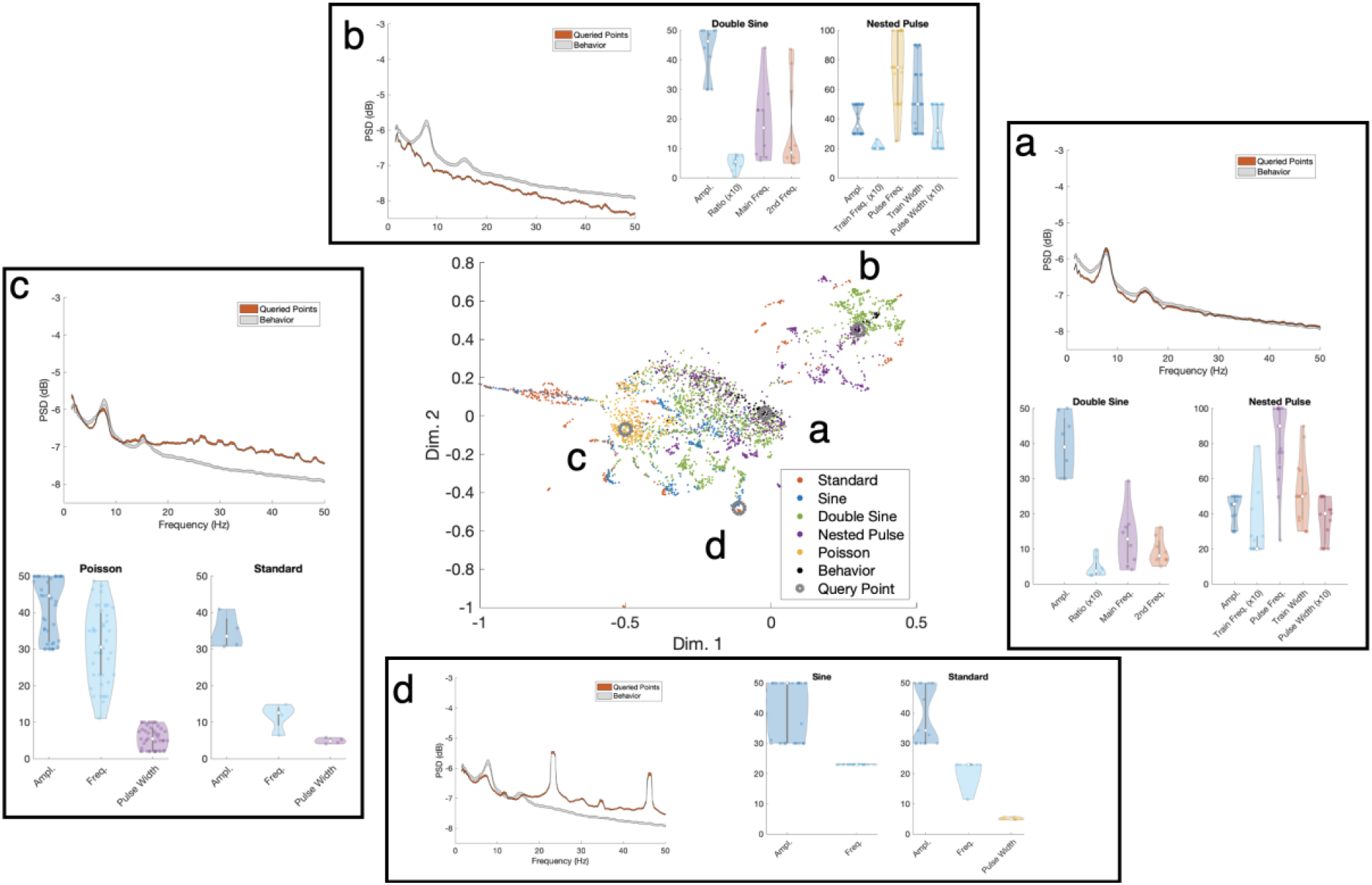
Inspection of the neural latent space. Recreated in the center is a representative plot of the neural latent space for one subject, with four manually selected query points shown as grey circles. For the respective points, each of panels a-d shows the PSD averaged for 50 data points nearest to the manually chosen point, the PSD averaged for all behavioral trials for comparison, and the values of stimulation parameters that were applied during the given trial. The total number of trials associated with each stimulation category is provided here: a) 26 behavioral trials; 15 nested pulse trials; 8 double sine trials; 1 sine trial. b) 6 behavioral trials, 10 double sine trials, 34 nested pulse trials. c) 44 Poisson trials; 4 standard pulse trials; 2 double sine trials. d) 15 standard pulse trials; 34 sine trials; 1 double sine trial. For double sine parameters, the “main frequency” is the frequency of the sinusoid that had a greater magnitude contribution to the summed waveform (i.e., a ratio parameter greater than 0.5). The values of some parameters are arbitrarily magnified by the factor shown in the x-axis labels so that their distributions are more clearly visible.

## 4. Discussion

There are several main conclusions of this study. First, we found that pan-neuronal medial septum optogenetic stimulation produced robust increases in hippocampal gamma power (3.1) but minimal change in theta power (3.2) regardless of differences in the stimulation waveforms. We then found that the influence of stimulation frequency on the power and shape of induced oscillations was consistent across waveforms. Second, we showed that meaningful differences between stimulation waveforms arise when studying their effects on hippocampal activity in the neural latent space and that the activity patterns induced by standard constant-frequency pulsatile stimulation are predominantly artificial (e.g. most different from endogenously recorded neural activity) (3.3). In contrast, the range of activity patterns induced by all other evaluated parameter spaces aligned more closely with activity observed during behavior. Third, the dimensionality reduction-based analysis strategy used can be applied to future work for investigating novel temporal stimulation patterns that produce desired physiological effects. Fourth, we provide a comprehensive characterization of the parameters for each of the waveforms in this study and their effects on hippocampal biomarkers.

When investigating the effects of stimulation waveforms on single-band hippocampal biomarkers, we found several counter-intuitive results: pan-neuronal optogenetic stimulation of the medial septum was unable to increase low-frequency activity above its baseline value across multiple parameter spaces. Further, varying the temporal pattern of stimulation did not produce a significant difference in the entrainment of either spectral biomarker. The former observation is particularly surprising given that the medial septum is known to be a predominant driver of the hippocampal theta rhythm.^10, 11^ Taken together, these results contradict common conventions in translational neuromodulation studies. For example, a study investigating electrical stimulation in the medial septum as a treatment for epilepsy might choose theta-frequency or theta-burst stimulation based on literature showing that hippocampal theta rhythms are seizure-resistant, then measure its effect on seizure frequency compared to sham stimulation (based on a mechanistic hypothesis of entraining hippocampal theta rhythms).^27, 28^ Given the finding that medial septum stimulation does not increase theta power, the fundamental mechanistic premise behind such a study may be incorrect but could easily go untested due to the challenges of measuring neural activity during electrical stimulation and evaluating its effect on behavioral symptoms. Ultimately, because the effects of brain stimulation can be counter-intuitive and could vary nonlinearly with the brain region and modality being studied, we propose that translational neuromodulation studies should always have an acute phase where the effects of multiple candidate stimulation patterns on neural activity are measured empirically to assert that underlying mechanistic assumptions are true before evaluating their therapeutic effect.

The analysis methods of our study could be applied to such translational studies with the following steps:

1. First, identify a natural neural signature that is associated with reduced disease symptoms.
2. Identify and test a wide array of candidate stimulation waveforms and parameters that may be well-suited to driving the desired neural activity signature.
3. Visualize the neural effects of each stimulation pattern in the neural latent space and quantify their similarity to the cluster of desired neural activity.
4. Identify the most promising subset of stimulation patterns for therapeutic testing.

Future work studying a wider array of neural circuits may extend our analysis methods to unravel whether our observed effects of different stimulation patterns are unique to optogenetic stimulation in the medial septum or might be fundamental to stimulation of neural systems and/or consistent with effects observed during equivalent temporal patterns of electrical stimulation. The extension of these methods to challenging electrical stimulation domains may be aided by recent well-validated advancements in artifact rejection^29^ or by measuring evoked potentials observed in-between stimulation pulses.^30^ The specific dimensionality reduction strategy could also be augmented by using novel methods that are targeted towards specific types of structure in neural data, including calcium imaging^31^ or the dynamics of neural spiking.^32^

One observation was that the higher-dimensional parameter spaces associated with the nested pulse and double sine waveforms did not produce a larger representation in the neural latent space than standard stimulation, let alone one that was quantitatively consistent with their increased dimensionality. This increased size comes at a cost, as the addition of each new parameter results in an exponential increase in the amount of data needed to understand and model the effects of stimulation or traverse the parameter space to find effective stimulation settings for a given application or subject.^7^ This “combinatorial explosion” effect is seen in the development of commercial deep brain stimulation devices, which have grown in complexity from thousands to millions of total parameter combinations available over years of clinical application.^33^ The addition of qualitatively novel temporal stimulation patterns may be a more effective strategy to expand the capability of neuromodulation devices, rather than adding new parameter dimensions which may be accompanied by much greater redundancy in their improved potential to modulate neural activity.

Out of the waveforms tested, we showed that conventional pulsatile stimulation induces patterns of neural activity that were most different from behavioral measurements. There are two specific features unique to standard pulse stimulation that may explain the artificiality of the corresponding induced neural activity: the abrupt rising edge of rectangular stimulation pulses and the constant-frequency nature of the waveform. All other waveforms examined in this study differed by either smoothly changing the amplitude over time (sinusoid and double sinusoid) and/or distributing the frequency content of the waveform into a mixture of slow and fast frequencies (nested pulse train and Poisson). Waveforms with these factors suppressed the high-magnitude harmonics of the stimulating frequency that were observed with standard pulse stimulation. Rectangular pulses might drive artificial neural responses by promoting highly synchronous firing across the spatial volume of tissue activated at the rising edge of the stimulation pulse, whereas a smooth increase in the amplitude of stimulation would result in different neuronal populations firing at different times as the spatial field of light, and therefore the corresponding volume of tissue activated, expands. Future work may investigate this hypothesis by examining the timing of neuronal population responses across a microelectrode array during pulsatile vs. sinusoidal stimulation and examine whether the biomimetic utility of sinusoidal stimulation can be replicated with electrical stimulation by smoothly adjusting the amplitude of an electrical pulse train.^34^

Trial-to-trial variability is a key limitation of the analysis in this study, which may result from multiple sources of state-dependence in the effects of stimulation.^35^ The dimensionality reduction strategy used was insufficient to precisely resolve differences between stimulation trials because the effects of stimulation nonlinearly summate with the animal’s natural neural activity state which varies throughout the experiment, and the modulation effect of stimulation itself at various frequencies can change nonlinearly based on the animal’s behavioral state.^16, 36^ Here we studied open-loop stimulation because it is more clinically feasible and there are major technical barriers to achieving precision state-dependent control of neural systems in patient care.^37^ Future work may explore how a dimensionality reduction approach can be augmented to identify the neural state components that predict variance in the effect of stimulation and achieve more precise modeling of stimulation effects. Such approaches may also be combined with novel data-driven optimization algorithms to search for subject-specific stimulation patterns that maximize the similarity between the stimulation-induced and desired brain state.^38^ Our work may also supplement the use of genetic algorithms and biophysically informed computational models of neural systems, which can suggest many more candidate stimulation patterns for empirical testing but suffer from limited ecological validity.^39^ Long-term, the integration of data-driven modeling and millisecond-timescale feedback control may be sufficient to precisely drive desired activity patterns in neural circuits to causally interrogate their function and treat disease.^40^

We found that harmonic entrainment of frequencies outside the stimulated frequency band was a key contributor to the artificial effects of standard pulse stimulation, but the physiological mechanisms of these harmonics remain incompletely understood. Various computational modeling results provide evidence that the recurrently connected structure of neuronal networks can give rise to both superharmonic and subharmonic entrainment effects when driven by external stimuli.^41, 42^ However, in experimental results these effects may partially or fully arise as an artifact of standardized Fourier transform-based analysis methods, which can identify harmonic power in signals that simply deviate from a perfectly sinusoidal waveform shape (but do not have a true physiological oscillation at the harmonic of the central frequency).^43^ Studying such harmonics in experimental recordings may be aided by novel analysis methods which have been developed to suppress such harmonics for the study of cross-frequency coupling in neural systems.^44^ A common neural control objective may be to drive a central neural oscillation without inducing harmonics in other oscillations which could be harmful; for example, Parkinsonian dyskinesia is associated with power in a subharmonic band of clinically standard DBS frequencies^45, 46^, and prior studies of DBS in epilepsy have been designed to avoid harmonics in frequencies that are expected to induce seizures.^47^ We observed that the Poisson waveform, when compared to standard pulse stimulation at an equivalent central frequency, increases power at the central frequency while obscuring the effects of harmonics. These empirical observations support recent theoretical work showing that adding noise to the inter-pulse interval of a constant-frequency stimulation waveform (“dithering”) can suppress harmonic stimulation effects.^48^ There may be a quantitative tradeoff between these two properties: having low inter-pulse noise produces greater entrainment at the central frequency, but at the cost of increased harmonics; higher inter-pulse noise might lead to reduced harmonics, but also decreased entrainment at the central frequency.

The finding that irregular temporal stimulation patterns can induce naturalistic neural activity patterns presents great utility for a variety of emerging neuropsychiatric conditions that can benefit from precision brain stimulation. Recent work in the treatment of depression led to significant symptom reduction when selecting patient-specific DBS parameters that produced a brain-wide electrophysiological network state that correlated with positive mood, and a strategy to improve memory was successful by driving the brain from “bad” into “good” information encoding states.^49, 50^ Various therapeutic applications have also benefited from identifying biomimetic stimulation patterns that are inspired by the characteristics of natural stimuli or the firing patterns of the population being targeted, including the restoration of touch via somatosensory cortical stimulation in brain-computer interfaces,^51^ the improvement of memory with hippocampal or amygdala stimulation^52, 53^, and visual cortex stimulation for restoring sight.^54^ The exploration of novel stimulation waveforms using scalable analysis strategies could benefit any of these applications by finding patterns that more accurately and consistently induce the desired neural activity state.

## REFERENCES

1. Lozano AM, Lipsman N, Bergman H, Brown P, Chabardes S, Chang JW, et al. Deep brain stimulation: current challenges and future directions Nat Rev Neurol. 2019 Mar;15:148–160.

2. Benabid AL. Deep brain stimulation for Parkinson’s disease Curr Opin Neurobiol. 2003 Dec;13:696–706.

3. Salanova V, Witt T, Worth R, Henry TR, Gross RE, Nazzaro JM, et al. Long-term efficacy and safety of thalamic stimulation for drug-resistant partial epilepsy Neurology. 2015 Mar 10;84:1017–1025.

4. Grill WM. Temporal Pattern of Electrical Stimulation is a New Dimension of Therapeutic Innovation Curr Opin Biomed Eng. 2018 Dec;8:1–6.

5. Brocker DT, Swan BD, So RQ, Turner DA, Gross RE, Grill WM. Optimized temporal pattern of brain stimulation designed by computational evolution Sci Transl Med. 2017 Jan 4;9.

6. Spix TA, Nanivadekar S, Toong N, Kaplow IM, Isett BR, Goksen Y, et al. Population-specific neuromodulation prolongs therapeutic benefits of deep brain stimulation Science. 2021 Oct 8;374:201–206.

7. Connolly MJ, Cole ER, Isbaine F, de Hemptinne C, Starr PA, Willie JT, et al. Multi-objective data-driven optimization for improving deep brain stimulation in Parkinson’s disease J Neural Eng. 2021 May 5;18.

8. Picillo M, Lozano AM, Kou N, Puppi Munhoz R, Fasano A. Programming Deep Brain Stimulation for Parkinson’s Disease: The Toronto Western Hospital Algorithms Brain Stimul. 2016 May-Jun;9:425–437.

9. Emiliani V, Cohen AE, Deisseroth K, Hausser M. All-Optical Interrogation of Neural Circuits J Neurosci. 2015 Oct 14;35:13917–13926.

10. Colgin LL. Rhythms of the hippocampal network Nat Rev Neurosci. 2016 Apr;17:239–249.

11. Cole ER, Grogan DP, Laxpati NG, Fernandez A, Skelton H, Isbaine F, et al. Evidence Supporting Deep Brain Stimulation of the Medial Septum in the Treatment of Temporal Lobe Epilepsy Epilepsia. 2022 Jun 14.

12. Laxpati NG, Mahmoudi B, Gutekunst CA, Newman JP, Zeller-Townson R, Gross RE. Real-time in vivo optogenetic neuromodulation and multielectrode electrophysiologic recording with NeuroRighter Front Neuroeng. 2014;7:40.

13. Lin JY, Lin MZ, Steinbach P, Tsien RY. Characterization of engineered channelrhodopsin variants with improved properties and kinetics Biophys J. 2009 Mar 4;96:1803–1814.

14. Park SE, Laxpati NG, Gutekunst CA, Connolly MJ, Tung J, Berglund K, et al. A Machine Learning Approach to Characterize the Modulation of the Hippocampal Rhythms Via Optogenetic Stimulation of the Medial Septum Int J Neural Syst. 2019 Dec;29:1950020.

15. Connolly MJ, Park SE, Laxpati NG, Zaidi SA, Ghetiya M, Fernandez A, et al. A framework for designing data-driven optimization systems for neural modulation J Neural Eng. 2020 Dec 3.

16. Mouchati PR, Kloc ML, Holmes GL, White SL, Barry JM. Optogenetic “low-theta” pacing of the septohippocampal circuit is sufficient for spatial goal finding and is influenced by behavioral state and cognitive demand Hippocampus. 2020 Nov;30:1167–1193.

17. Bokil H, Andrews P, Kulkarni JE, Mehta S, Mitra PP. Chronux: a platform for analyzing neural signals J Neurosci Methods. 2010 Sep 30;192:146–151.

18. Rasmussen CE, Nickisch H. Gaussian processes for machine learning (GPML) toolbox The Journal of Machine Learning Research. 2010;11:3011–3015.

19. McInnes L, Healy J, Melville J. UMAP: Uniform Manifold Approximation and Projection for Dimension Reduction2018 February 01, 2018:[arXiv:1802.03426 p.]. Available from: https://ui.adsabs.harvard.edu/abs/2018arXiv180203426M.

20. Lee EK, Balasubramanian H, Tsolias A, Anakwe SU, Medalla M, Shenoy KV, et al. Non-linear dimensionality reduction on extracellular waveforms reveals cell type diversity in premotor cortex Elife. 2021 Aug 6;10.

21. Van der Maaten L, Hinton G. Visualizing data using t-SNE Journal of machine learning research. 2008;9.

22. Kobak D, Berens P. The art of using t-SNE for single-cell transcriptomics Nat Commun. 2019 Nov 28;10:5416.

23. Shlens J. A Tutorial on Principal Component Analysis2014 April 01, 2014:[arXiv:1404.1100 p.]. Available from: https://ui.adsabs.harvard.edu/abs/2014arXiv1404.1100S.

24. Peter D H. Kernel estimation of a distribution function Communications in Statistics - Theory and Methods. 2007;14:605–620.

25. Dice LR. Measures of the Amount of Ecologic Association Between Species Ecology. 1945;26:297–302.

26. Cole ER, Grogan DP, Eggers TE, Connolly MJ, Laxpati NG, Gross RE. Model-Driven Collection of Neural Modulation Data. 2021 10th International IEEE/EMBS Conference on Neural Engineering (NER)2021. p. 281–284.

27. Lee DJ, Izadi A, Melnik M, Seidl S, Echeverri A, Shahlaie K, et al. Stimulation of the medial septum improves performance in spatial learning following pilocarpine-induced status epilepticus Epilepsy Res. 2017 Feb;130:53–63.

28. Park SE, Connolly MJ, Exarchos I, Fernandez A, Ghetiya M, Gutekunst CA, et al. Optimizing neuromodulation based on surrogate neural states for seizure suppression in a rat temporal lobe epilepsy model J Neural Eng. 2020 Jul 16;17:046009.

29. Dastin-van Rijn Em, Provenza NR, Calvert JS, Gilron R, Allawala AB, Darie R, et al. Uncovering biomarkers during therapeutic neuromodulation with PARRM: Period-based Artifact Reconstruction and Removal Method Cell Rep Methods. 2021 Jun 21;1.

30. Dale J, Schmidt SL, Mitchell K, Turner DA, Grill WM. Evoked potentials generated by deep brain stimulation for Parkinson’s disease Brain Stimul. 2022 Jul 31.

31. Saxena S, Kinsella I, Musall S, Kim SH, Meszaros J, Thibodeaux DN, et al. Localized semi-nonnegative matrix factorization (LocaNMF) of widefield calcium imaging data PLoS Comput Biol. 2020 Apr;16:e1007791.

32. Pandarinath C, O’Shea DJ, Collins J, Jozefowicz R, Stavisky SD, Kao JC, et al. Inferring single-trial neural population dynamics using sequential auto-encoders Nat Methods. 2018 Oct;15:805–815.

33. Steigerwald F, Matthies C, Volkmann J. Directional Deep Brain Stimulation Neurotherapeutics. 2019 Jan;16:100–104.

34. Tan DW, Schiefer MA, Keith MW, Anderson JR, Tyler J, Tyler DJ. A neural interface provides long-term stable natural touch perception Sci Transl Med. 2014 Oct 8;6:257ra138.

35. Bradley C, Nydam AS, Dux PE, Mattingley JB. State-dependent effects of neural stimulation on brain function and cognition Nat Rev Neurosci. 2022 Aug;23:459–475.

36. Cole ER, Connolly MJ, Park S-E, Grogan DP, Buxton W, Eggers TE, et al. Autonomous State Inference for Data-Driven Optimization of Neural Modulation. 2021 10th International IEEE/EMBS Conference on Neural Engineering (NER)2021. p. 950–953.

37. Arlotti M, Rosa M, Marceglia S, Barbieri S, Priori A. The adaptive deep brain stimulation challenge Parkinsonism Relat Disord. 2016 Jul;28:12–17.

38. Schrum M, Connolly MJ, Cole E, Ghetiya M, Gross R, Gombolay MC. Meta-Active Learning in Probabilistically Safe Optimization IEEE Robotics and Automation Letters. 2022:1–8.

39. Cassar IR, Titus ND, Grill WM. An improved genetic algorithm for designing optimal temporal patterns of neural stimulation J Neural Eng. 2017 Dec;14:066013.

40. Bolus MF, Willats AA, Rozell CJ, Stanley GB. State-space optimal feedback control of optogenetically driven neural activity J Neural Eng. 2021 Mar 31;18.

41. Velarde OM, Urdapilleta E, Mato G, Dellavale D. Bifurcation structure determines different phase-amplitude coupling patterns in the activity of biologically plausible neural networks Neuroimage. 2019 Nov 15;202:116031.

42. Roberts JA, Robinson PA. Quantitative theory of driven nonlinear brain dynamics Neuroimage. 2012 Sep;62:1947–1955.

43. Dellavale D, Velarde OM, Mato G, Urdapilleta E. Complex interplay between spectral harmonicity and different types of cross-frequency couplings in nonlinear oscillators and biologically plausible neural network models Phys Rev E. 2020 Dec;102:062401.

44. Dellavale D, Urdapilleta E, Campora N, Velarde OM, Kochen S, Mato G. Two types of ictal phase-amplitude couplings in epilepsy patients revealed by spectral harmonicity of intracerebral EEG recordings Clin Neurophysiol. 2020 Aug;131:1866–1885.

45. Swann NC, de Hemptinne C, Miocinovic S, Qasim S, Wang SS, Ziman N, et al. Gamma Oscillations in the Hyperkinetic State Detected with Chronic Human Brain Recordings in Parkinson’s Disease J Neurosci. 2016 Jun 15;36:6445–6458.

46. Sermon JJ, Olaru M, Anso J, Little S, Bogacz R, Starr PA, et al. Sub-harmonic Entrainment of Cortical Gamma Oscillations to Deep Brain Stimulation in Parkinson’s Disease: Predictions and Validation of a Patient-Specific Nonlinear Model bioRxiv. 2022:2022.2003.2001.482549.

47. Zamora M, Meller S, Kajin F, Sermon JJ, Toth R, Benjaber M, et al. Case Report: Embedding “Digital Chronotherapy” Into Medical Devices-A Canine Validation for Controlling Status Epilepticus Through Multi-Scale Rhythmic Brain Stimulation Front Neurosci. 2021;15:734265.

48. Duchet B, Sermon JJ, Weerasinghe G, Denison T, Bogacz R. How to entrain a selected neuronal rhythm but not others: open-loop dithered brain stimulation for selective entrainment bioRxiv. 2022:2022.2007.2006.499051.

49. Sheth SA, Bijanki KR, Metzger B, Allawala A, Pirtle V, Adkinson JA, et al. Deep Brain Stimulation for Depression Informed by Intracranial Recordings Biol Psychiatry. 2021 Nov 22.

50. Ezzyat Y, Wanda PA, Levy DF, Kadel A, Aka A, Pedisich I, et al. Closed-loop stimulation of temporal cortex rescues functional networks and improves memory Nat Commun. 2018 Feb 6;9:365.

51. Lee B, Kramer D, Armenta Salas M, Kellis S, Brown D, Dobreva T, et al. Engineering Artificial Somatosensation Through Cortical Stimulation in Humans Front Syst Neurosci. 2018;12:24.

52. Hampson RE, Song D, Robinson BS, Fetterhoff D, Dakos AS, Roeder BM, et al. Developing a hippocampal neural prosthetic to facilitate human memory encoding and recall J Neural Eng. 2018 Jun;15:036014.

53. Inman CS, Manns JR, Bijanki KR, Bass DI, Hamann S, Drane DL, et al. Direct electrical stimulation of the amygdala enhances declarative memory in humans Proc Natl Acad Sci U S A. 2018 Jan 2;115:98–103.

54. Beauchamp MS, Oswalt D, Sun P, Foster BL, Magnotti JF, Niketeghad S, et al. Dynamic Stimulation of Visual Cortex Produces Form Vision in Sighted and Blind Humans Cell. 2020 May 14;181:774–783 e775.

