## Supplemental Figures for "Irregular optogenetic stimulation waveforms can induce naturalistic patterns of hippocampal spectral activity"

### SUPPLEMENTARY MATERIAL

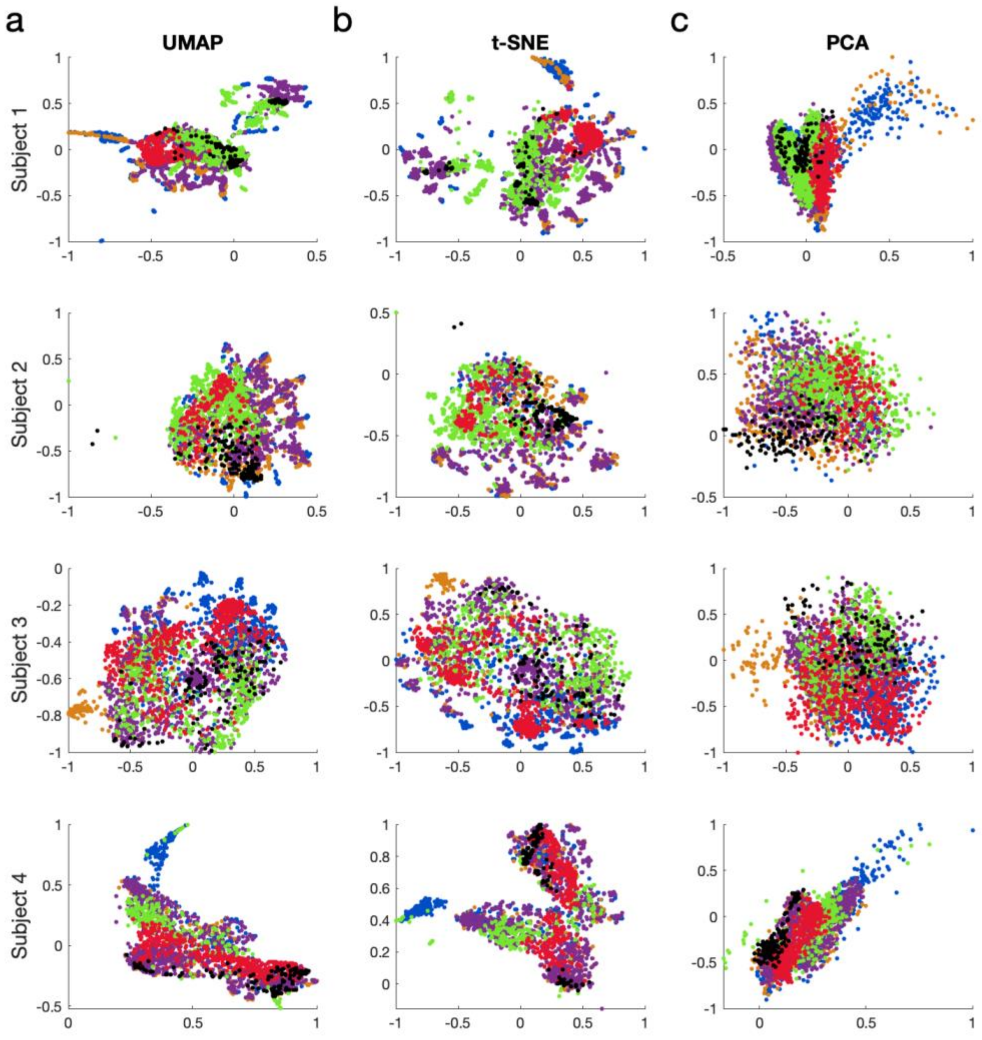


Figure S1: All subject-specific latent space representations for all combinations of four subjects and 3 dimensionality reduction strategies.

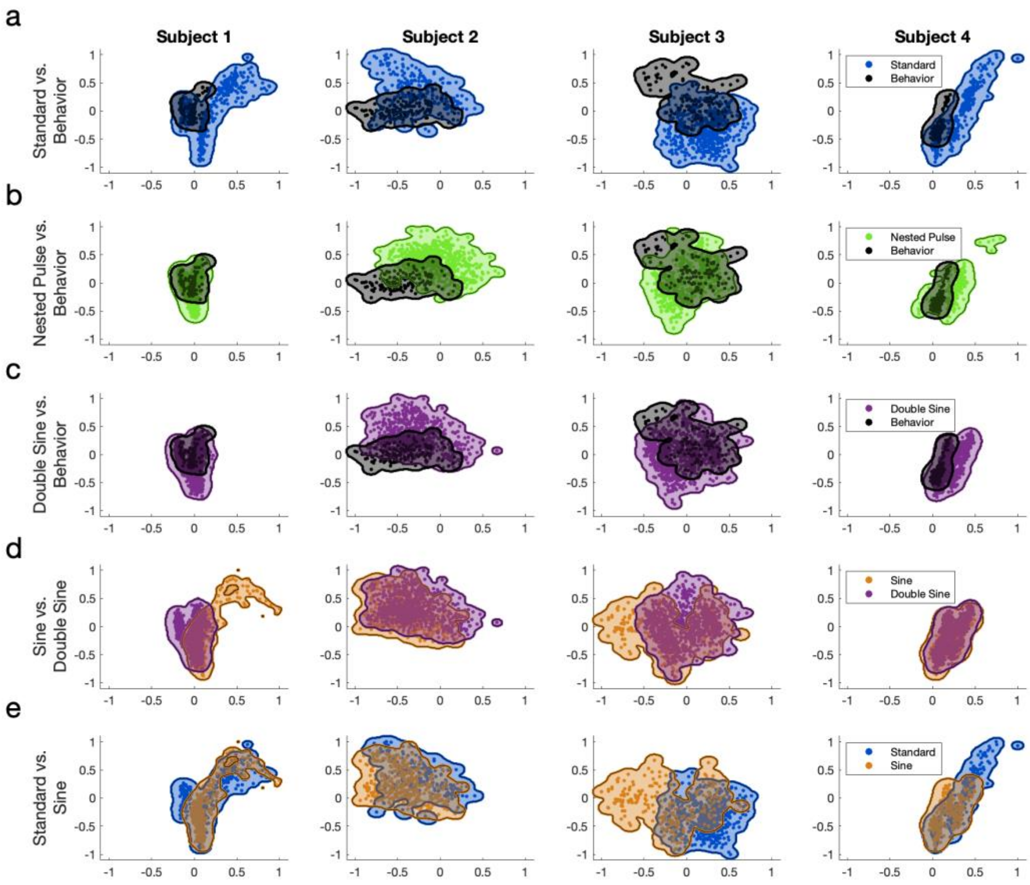


Figure S2: Subject-specific pairwise comparisons of parameter space latent representations.

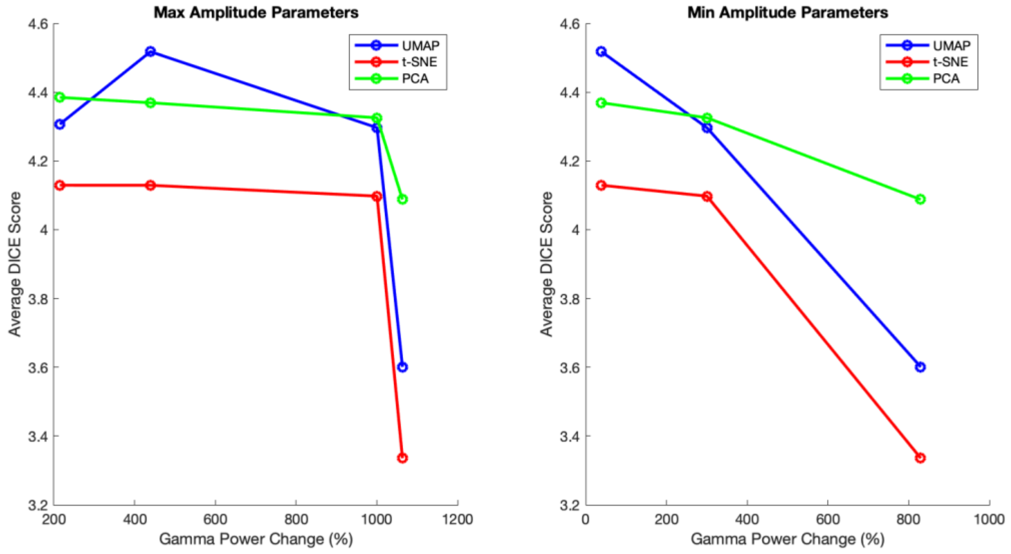


Figure S3: Subject-specific variability in the "cloudiness" of DR representations is explained by the magnitude of their optogenetic stimulation response. Y-axis: Magnitude of the average overlap between all parameter spaces, plotted separately for 4 subjects and 3 dimensionality reduction algorithms. X-axis: The magnitude of power increase from baseline observed for both maximum (left) and minimum (right) amplitude stimulation parameters. The negative trend in average overlap vs. strength of the optogenetic response (especially with the minimum amplitude parameters) provides an explanation for the between-subject variability in the separability of the latent space representations: less clearly separable representations may result from the weaker effects of lower-amplitude stimulation parameters, whereas the most clearly separable representation was observed for the subject with the strongest response at low-amplitude parameters.

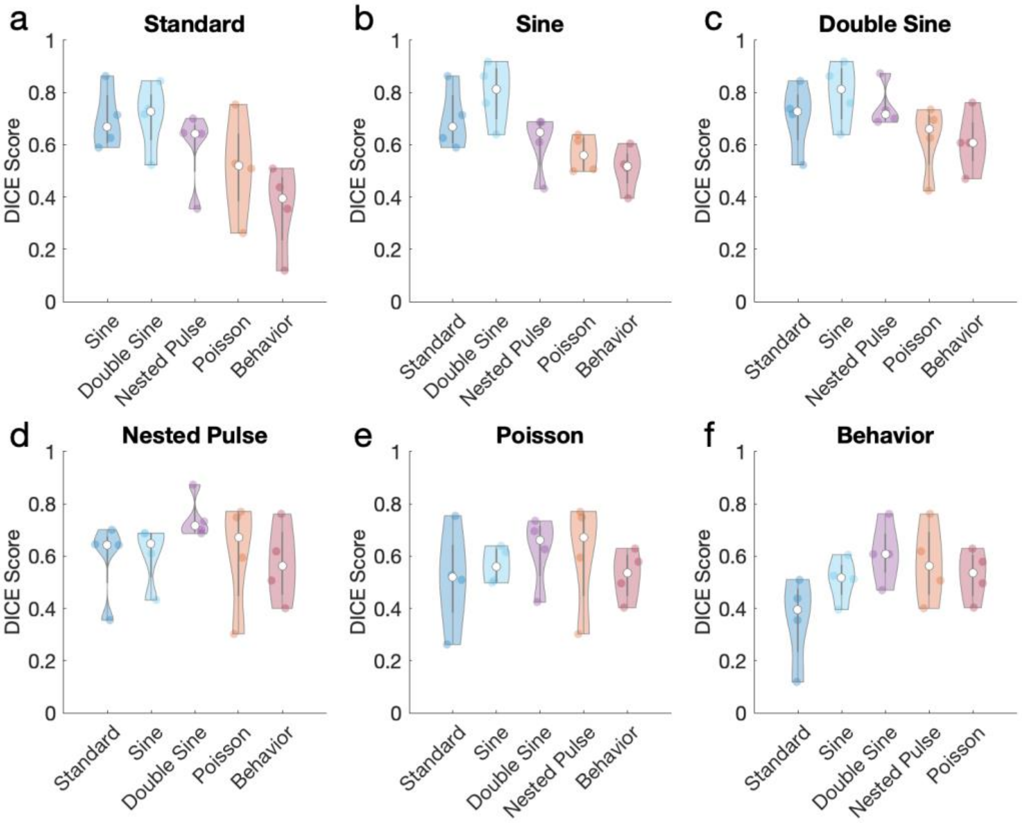


Figure S4: Complete set of pairwise DICE score comparisons across all parameter spaces.

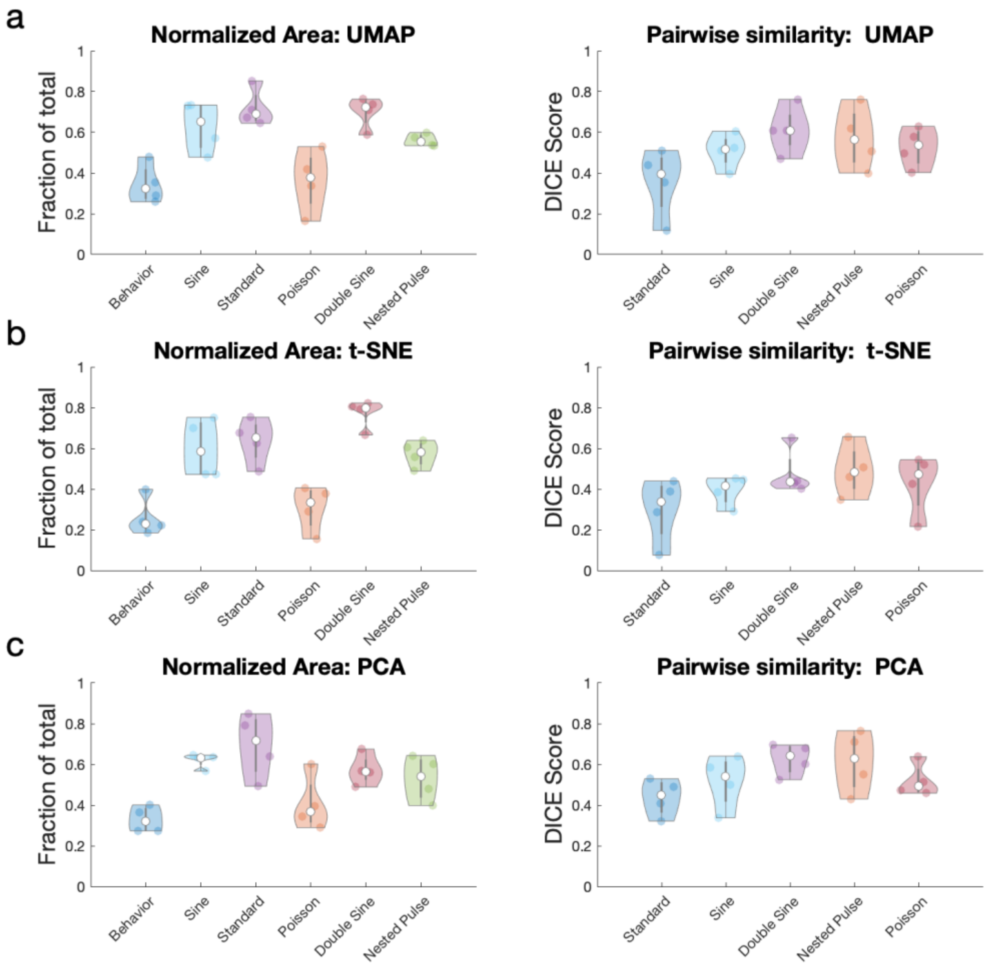


Figure S5: Figure panels 4c and 4f from the main text are replicated for all three dimensionality algorithms used in the study (a: UMAP, shown in figure 4; b: t-SNE; and c: PCA). The trends observed in part a are replicated in both b and c.
